## Supplementary material for "An Evolutionary Process Without Variation and Selection": Alphabetical list of Definitions of Key Terms

LIANE GABORA AND MIKE STEEL

Addresses for Correspondence:

Liane Gabora
Department of Psychology, University of British Columbia, Kelowna BC, Canada


Mike Steel
Biomathematics Research Centre, University of Canterbury, Christchurch, New Zealand


### 1. Alphabetical list of Definitions of Key Terms

Since miscommunication arises from inconsistencies in how evolutionary concepts are applied in the sciences and social sciences, to maintain clarity we provide an alphabetical list of standard definitions of key terms used in this paper.

- **Acquired trait:** a trait obtained during the lifetime of its bearer (e.g., a scar, tattoo, or the memory of a song) and transmitted horizontally (i.e., laterally).
- **Culture:** extrasomatic adaptations—including behavior and artifacts—that are socially rather than sexually transmitted.
- **Darwinian (or 'selectionist') process:** an evolutionary process—i.e., a process that exhibits cumulative, adaptive, open-ended change—occurring through natural or artificial selection.
- **Darwinian threshold:** transition from a non-Darwinian to a Darwinian evolutionary process [13, 12].
- **Evolution:** descent with modification, or cumulative, adaptive change over time, giving rise to new species that share a common ancestor [3]. The most well-known theories to explain how evolution works are natural selection and Lamarckism. The concept of evolution has been extended to apply to, not just to cumulative, adaptive change in biological species, but also cumulative, adaptive *cultural* change [1, 2]. Thus, here we use the more general definition of evolution used in cultural evolution research: cumulative, adaptive change over time.
- **Generation:** a single transition period from the internalized to the externalized form of a trait.[1]

[1]Note that, with respect to biological evolution, a new generation generally (though not in horizontal gene transfer) begins with the birth of one or more organism(s). With respect to cultural evolution, a 'generation' may begin with the transmission of an idea (a cultural trait). Thus, over the course of a discussion, the idea may undergo multiple generations.

- **Horizontal transmission:** non-germ-line transmission of an acquired trait from one entity to another. Thus, social transmission is horizontal because the information is not inherited, i.e., it is not passed from parent to offspring by way of genes.
- **Lamarckian process (or 'soft inheritance'):** widely understood to mean the transmission of characteristics to offspring that were acquired through use or disuse during its lifetime.[2] Some maintain that Lamarckian evolution requires genetic transmission to biological offspring, a view held by early biologists (e.g., [10]), though current scholars (e.g., [9]) sometimes take a more equivocal view.
- **Neutral theory of molecular evolution**: the theory that a significant degree of genetic variation in populations is the result of mutation and genetic drift, not selection [6, 7].
- **Selection:** differential replication of randomly generated heritable variation in a population over generations such that some traits become more prevalent than others. Selection may be natural (due to non-human elements of the environment) or artificial (due to human efforts such as selective breeding), and it can occur at multiple levels, e.g., genes, individuals, or groups [8].
- **Self-assembly code:** a set of coded instructions that is: (i) *actively interpreted* through a developmental process to generate a soma, and (ii) *passively copied without interpretation* through a reproduction process to generate self-copies, i.e., new sets of self-assembly instructions that are in turn used in these two distinct ways. (The word 'instructions' should not be interpreted literally here as to imply that they are written in a natural language such as English. The instructions are implicit in the dynamics of the entity; it is structured in such a way as to ensure a sequence of processes that result in self-assembly.)

[2]We note that although this is how the term is standardly defined, it has been pointed out that this idea was not unique to Lamarck, and transmission of characteristics was only one component of Lamarck's theory [5].

- **Self–Other Reorganisation (SOR):** a theory of how both culture, and early life, evolve through interacting, self-organizing networks, based on theory and findings that have come out of the post-modern synthesis in biology [4, 11].
- **Vertical transmission:** Germ-line inheritance of a trait from one generation to the next. (In other words, transmission occurs by way of a self-assembly code, such as DNA.)
